# Bidirectional redistribution of actomyosin drives epithelial invagination in ascidian siphon tube morphogenesis

**DOI:** 10.1101/2025.08.26.672310

**Authors:** Jinghan Qiao, Pengyu Yu, Hongzhe Peng, Wenjie Shi, Bo Li, Bo Dong

## Abstract

How epithelia perform a spatiotemporal heterogeneous force generating program to drive a sequential tissue morphogenesis remains unclear, particularly the underlying precise mechanical mechanisms. This study investigated dynamic actomyosin reorganization between apical and lateral membrane cortex regions during two sequentially invaginated stages during ascidian atrial siphon tube morphogenesis. At the initial invagination stage, the originally lateral-located actomyosin redistributed to the apical domains, while those actomyosin redistributed back to lateral domains at the accelerated invagination stage. Using genetic mutants to modulate myosin activities, the initial invagination was strengthened or abolished, indicating invagination is apical constriction dependent. Optogenetic inhibition of myosin activities in lateral domains after initial invagination stage blocked the further processes, suggesting lateral constriction of actomyosin is required for the accelerated invagination. Vertex model simulations uncovered a coupled mechanism underlying epithelial invagination driven by apicobasal tension imbalance and lateral contraction. We thus propose an actomyosin redistribution mechanical model: lateral actomyosin first redistributes apically to drive apical constriction and shape the initial invagination, then apical actomyosin redistributes laterally to promote lateral contractility and accelerate invagination. Our findings reveal a bidirectional reorganization of actomyosin network as a central mechanism driving epithelial invagination, providing insights on epithelial invagination and the organ morphogenesis during development.

## Introduction

Morphogenesis, by which cells and tissues acquire their shape and structure, is a central theme in developmental biology. The tissue shape transformation in morphogenesis is fundamentally driven by the spatiotemporally regulated biomechanical forces(Collinet & Lecuit, 2021; Matter & Balda, 2003; Štorgel et al., 2016). The generation and conduction of biomechanical forces play as an essential mechanism of morphogenesis, via regulating cell rearrangement, division, and deformation at both cellular and tissue scales(Chan et al., 2017; Lecuit et al., 2011; Matter & Balda, 2003; Nagafuchi, 2001; Osswald et al., 2022; Tepass, 2002). Among these, actomyosin contractility plays a pivotal structure generating biomechanical tension in subcellular level, which is the basic of cell and tissue-level morphogenesis(Chung et al., 2017; Even-Ram et al., 2007; Fristrom, 1988; Lee & Harland, 2007; Popkova et al., 2024; Vicente-Manzanares et al., 2009; Zartman & Shvartsman, 2010). Despite the widely acknowledged critical role of bio-mechanical forces in morphogenesis, the precise mechanisms by which these forces regulate cell and tissue remodeling, particularly how cells reorganize contractile domains in space and time to coordinate sequential morphogenetic events remains a significant challenge.

A typical example of morphogenesis is the transition from simple epithelial sheets to complex three-dimensional architectures(Collinet & Lecuit, 2021). A large number of previous researches demonstrated the function of apical constriction in epithelial sheet invagination, such as in the invagination process of the *Drosophila* embryonic trachea(Schottenfeld et al., 2010) and salivary gland placode(Pearl et al., 2017). However, more and more studies noticed that the apical constriction itself is not sufficient in shaping complex epithelial three-dimensional structures and various epithelial tube shapes(Kondo & Hayashi, 2013; Pearl et al., 2017). So, an interesting question is how epithelial cells perform a spatiotemporal heterogeneous force generating program to drive a sequential epithelial tissue reshaping.

Marine ascidian *Ciona* is an excellent model for studying morphogenesis due to its simple developmental process, transparent embryos, and its close evolutionary relationship with vertebrates(Blair & Hedges, 2005; Delsuc et al., 2006; Zhao et al., 2021). For example, asymmetrical actomyosin contractility in notochord was demonstrated to provide force for tail bending(Lu et al., 2020; Peng et al., 2020). Local myosin activation coupled with junctional rearrangements drives directional zippering(Fiuza & Lemaire, 2021; Hotta et al., 2007; Kourakis et al., 2010) in neural tube closure, and cell cortex distribution and the stability of tight junctions (TJs) were essential for notochord tube lumen opening and expansion(Shi et al., 2025). Actomyosin sequentially localized on the apical and basolateral cell surfaces to drive the endodermal invagination(Fiuza & Lemaire, 2021; Hotta et al., 2007; Kourakis et al., 2010). However, this transition is from apical to basolateral(Sherrard et al., 2010), and the lateral domain does not play a significant role in this process. Similarly, in *Drosophila* gastrulation, apical constriction initiates ventral furrow formation, but lateral myosin is not absolutely required for the later stages; indeed, the second phase of invagination can proceed even when lateral contractility is compromised(Guo et al., 2022). This again suggests that the role of lateral contractility during invagination remains underexplored.

*Ciona* and vertebrates share evolutionary developmental homology. There may be a potential homology between the otic placode of vertebrates and the atrial siphon primordium of *Ciona*(Kourakis et al., 2010). The atrial siphon forms from a non-dividing region in the lateral-dorsal epidermis of the head and follows a separate developmental trajectory(Hotta et al., 2020; Kourakis et al., 2010). Invagination initially generates shallow pits, which later deepen and connect to the gut lumen after metamorphosis, creating two atrial siphons(Hotta et al., 2007; Kourakis et al., 2010). Eventually, the left and right siphons fuse to form a single atrial siphon(Chiba et al., 2004). Despite these observations, the early cellular events during atrial siphon tube morphogenesis remain incompletely understood.

In this study, we investigated the morphogenesis of the *Ciona* atrial siphon. Firstly, the detailed cellular process of *Ciona* atrial siphon formation through initial and the accelerated stages was described by visualizing cell boundary and nuclei. By combining actomyosin localization analysis, genetic manipulation, and vertex model simulations, we demonstrated that lateral actomyosin first redistributed apically to drive apical constriction and established the initial invagination, then redistributed laterally to promote cell shortening and the further atrial siphon tube invagination. Disruption of myosin activity altered invagination timing, while optogenetic inhibition of myosin activity after initial apical constriction stage blocked the further invagination. Furthermore, a cell-based vertex model was established to validate the relationship between contractile force redistribution and epithelial invagination. The coupled biomechanical mechanisms induced by apicobasal imbalance and lateral contraction were uncovered as fundamental determinants of the atrial siphon invagination. These results provide new insights into the biomechanical control of tissue morphogenesis, and highlight the value of marine model organisms for addressing fundamental questions in developmental biology.

## Results

### The cellular processes of *Ciona* atrial siphon invagination

To investigate the early cellular events during atrial siphon morphogenesis, we performed actin filament (F-actin) staining on the fixed samples to capture the detailed cellular dynamics (Figure 1A). The cell at the bottom center of the invaginating region was defined as the “center cell” (Figure 1B). Measurements taken at different stages included the height and apical-to-basal area ratio of the center cell, the invagination depth of the atrial siphon and the linear distance between the -3/-4 and +3/+4 cell junctions (Figure 1B-E, Figure 1—figure supplement 1). At the initial invagination stage (13.5-16.0 hpf), the center cell height increased, the apical-to-basal area ratio decreased, and the protruding cell surface became inward pit, but no significant invagination occurred (Figure 1A), as reflected by a low invagination slope (k = 0.2617). During the accelerated stage (16.0-18.0 hpf), the center cell height decreased, the apical-to-basal area ratio stabilized, and invagination progressed rapidly (k = 2.7920). EdU incorporation and TUNEL assays demonstrated that neither cell division nor apoptosis was involved in the whole invagination process (Figure 1—figure supplement 2).

**Figure 1.**
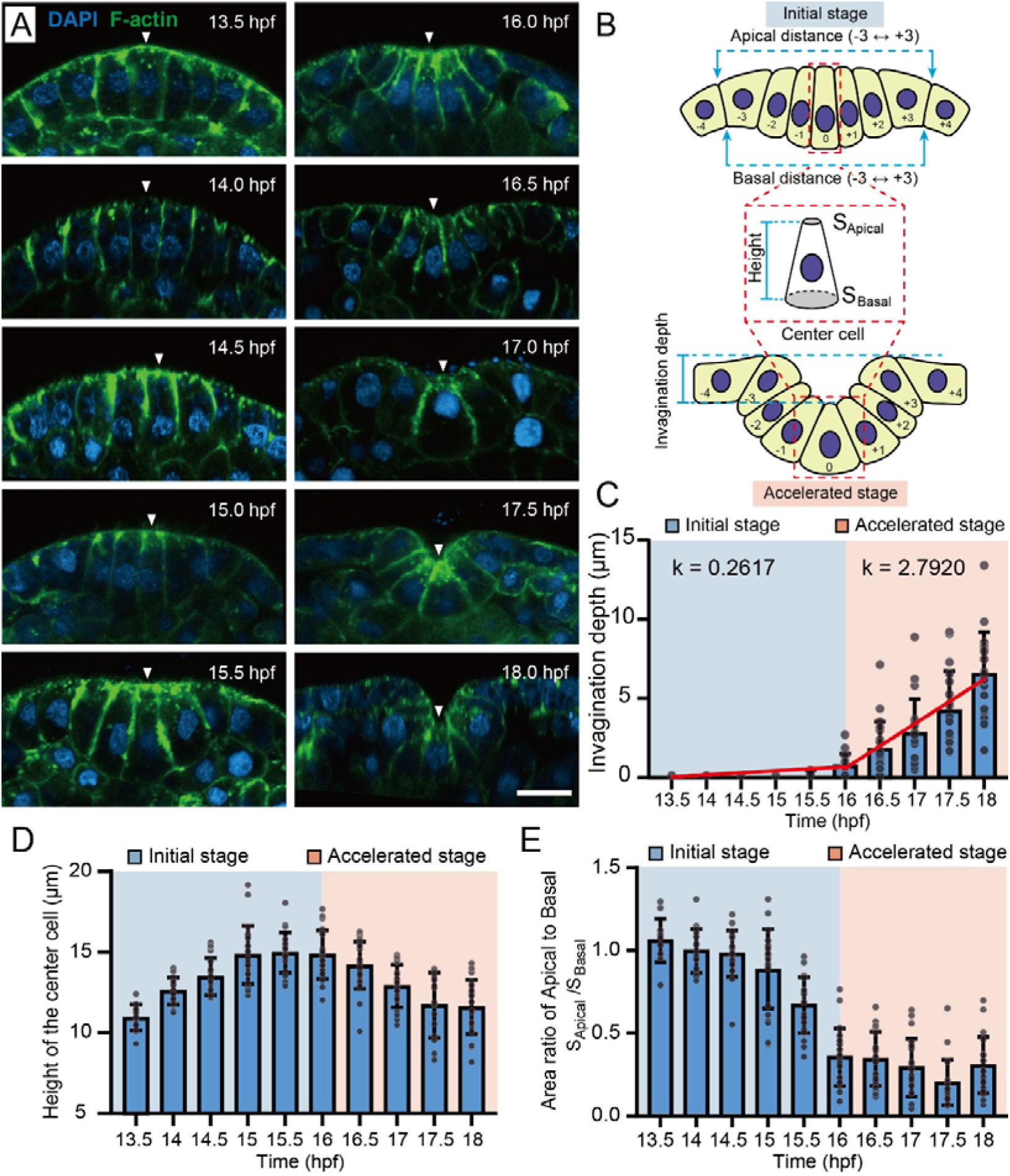
Cellular processes of *Ciona* atrial siphon tube invagination. (A) Representative images of atrial siphon morphogenesis in *Ciona* embryos from 13.5 hpf to 18 hpf. The white arrow indicates the central cell. Scale bar: 10 μm. (B) Measurement parameters of the *Ciona* atrial siphon. The cell undergoing the most prominent apical constriction at the center of invagination was defined as the center cell (0). The adjacent cells on the left and right were defined sequentially as -1, -2, -3, -4 and +1, +2, +3, +4, respectively. (C) Quantification of the invagination depth in the atrial siphon of *Ciona* embryos. Red lines indicate linear regression fits of invagination depth during the initial (13.5-16 hpf, k = 0.2617) and accelerated (16-18 hpf, k = 2.7920) stages. n = 20. (D, E) Quantification of the center cell height and apical-to-basal area ratio at the atrial siphon of *Ciona* embryos. The blue-shaded region represents the initial stage, while the orange-shaded region indicates the accelerated stage. n = 20.

### Bidirectional redistribution of actomyosin between apical and lateral domains during atrial siphon tube invagination

To analyze the forces driving atrial siphon tube invagination, we first visualized the distribution of F-actin and quantified its dynamics at different cellular domains (Figure 1—figure supplement 1A). During the initial invagination stage, before 15 hpf, F-actin concentration decreased at the lateral domains, while gradually increasing around the apical membrane, resulting in a higher F-actin accumulation at the apical region. Subsequently, the F-actin concentration at lateral region started to increase and exceeded that in the apical domain at 16 hpf (Figure 1—figure supplement 1A).

Using anti-pS19 MRLC antibody, we examined the spatial and temporal patterns of myosin activity within the atrial siphon primordium. Quantification of signal intensities from the apical, basal, and lateral regions during the initial and the accelerated stages showed that active myosin underwent a clear bidirectional redistribution (Figure 2A, B). During the initial invagination stage (before 16 hpf), myosin activity shifted from the lateral to the apical domain, resulting in higher apical levels. In contrast, during the accelerated stage (after 16 hpf), myosin redistributed from the apical to the lateral domain, leading to predominant lateral enrichment (Figure 2B). This redistribution pattern of active myosin showed a similar trend to that of F-actin in the corresponding phases, although the changes in F-actin did not reach statistical significance during the initial stage.

**Figure 2.**
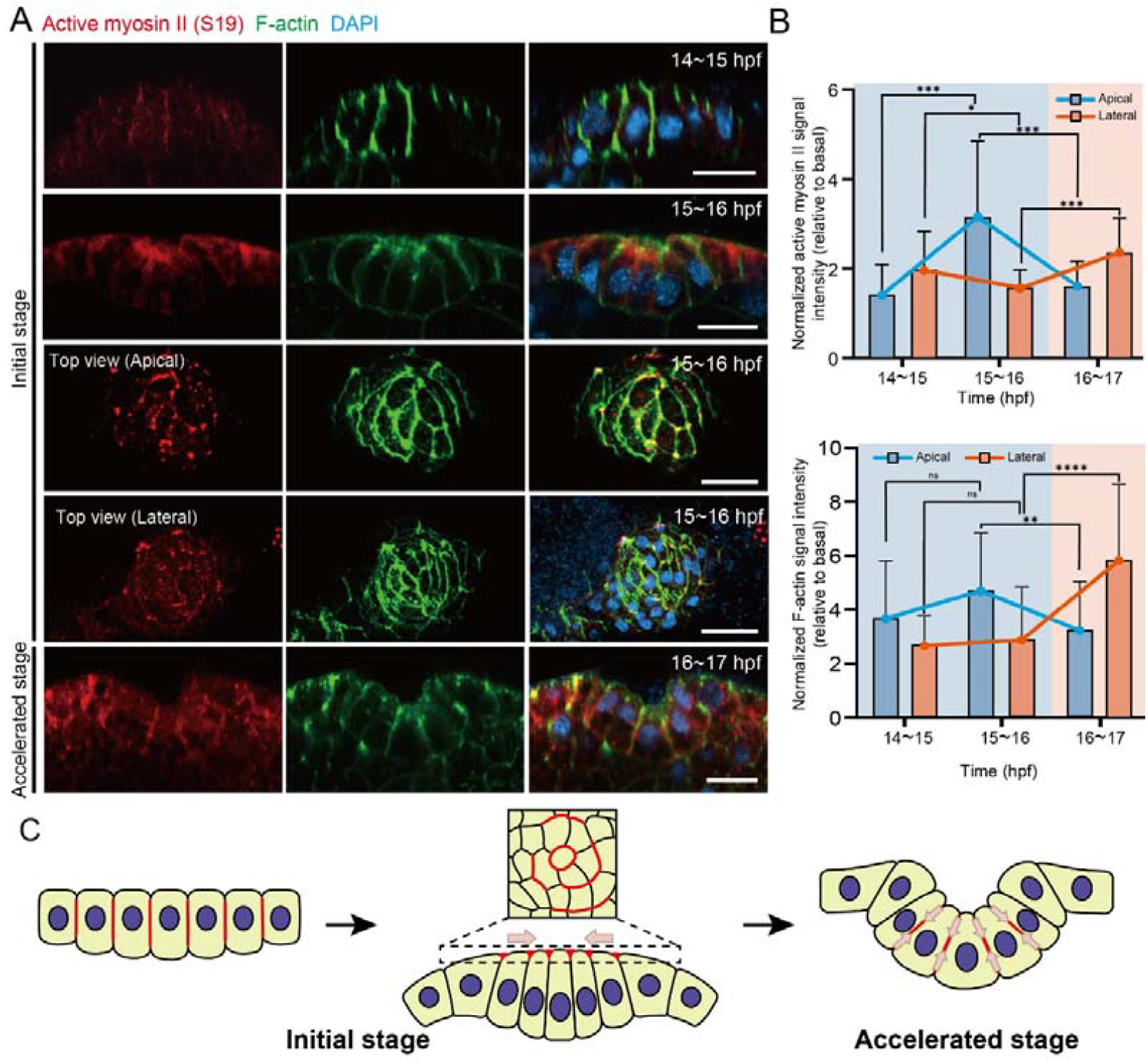
Bidirectional redistribution of actomyosin between apical and lateral domains during atrial siphon tube invagination. (A) Representative images of *Ciona* embryos stained for active myosin II (anti-pS19 MRLC,red) and F-actin (green) at different stages of atrial siphon invagination. Scale bar: 10 μm. (B) Normalized fluorescence intensity of active myosin II (anti-pS19 MRLC) and F-actin at the apical and lateral regions of center cells during different stages of atrial siphon invagination (basal level set to 1). The blue-shaded region represents the initial stage, while the orange-shaded region indicates the accelerated stage. **p < 0.01, ***p < 0.001, ****p < 0.0001, n = 20. (C) Schematic model illustrating the mechanical forces driving atrial siphon primordium invagination. Brown arrows indicate the direction of contractile forces; red areas depict active myosin II localization.

Additionally, top view imaging showed that during the initial stage, atrial siphon primordium cells were arranged in a circular pattern (Figure 2A). Active myosin exhibited a similar ring-like localization, and this ring-like pattern extended to about 15–20 cells of the primordium, indicating that the early stages of atrial siphon morphogenesis are driven by circumferential contractile forces generated by phosphorylated myosin, facilitating inward constriction of the primordium (Figure 2A, B). Furthermore, quantification of the lateral cell distance (Figure 1—figure supplement 1B) demonstrated a reduction in spacing between peripheral cells, indicating that surrounding cells moved centripetally toward the center position.

Integrating the localization of actomyosin with the observed cell shape changes, we propose a hypothesis in which the bidirectional reorganization of contractile forces at the apical and lateral regions drives epithelial invagination during atrial siphon morphogenesis (Figure 2C). Initially, actomyosin redistributed from the lateral regions to the apical domains, generating contractile forces for apical constriction (Figure 1—figure supplement 1A). Meanwhile, the inward compression from surrounding cells facilitates cell elongation, establishing the initial invaginated cell morphology and preparing for subsequent morphogenesis (Figure 1—figure supplement 1B). Then, actomyosin redistributed from the apical domains to the lateral regions, generating contractile forces that promoted center cell shortening and accelerated the tissue invagination (Figure 2C).

### Center cell height is coupled with invagination depth

To further verify our hypothesis and explore the relationship between myosin contractility, force redistribution, and the progression of invagination, we overexpressed MRLC (T18ES19E)::mCherry (a diphosphorylated mutant that enhances myosin activity) (Espinoza-Fonseca et al., 2014), MRLC (T18AS19A)::mCherry (an unphosphorylated mutant that reduces myosin activity) (Iwasaki et al., 2001) and MRLC::mCherry (wild-type control) in the atrial siphon primordium to modulate contractile force in embryos, respectively (Figure 3A, B). The results showed that the invagination initiated earlier (15 hpf) compared to control group in MRLC (T18ES19E)-expressed embryos. At 16 hpf, MRLC (T18ES19E) group exhibited a deeper invagination and shorter center cell height. In contrast, in MRLC (T18AS19A) group, the invagination initiation time was delayed. Even at 16 hpf, the center cell height did not decrease and the invagination did not occur (Figure 3B). Notably, the increase in invagination depth is strongly correlated with the reduction in center cell height, with both changes occurring synchronously across experimental groups with different myosin activities (Figure 3B). These results demonstrate a strong coupling between the reduction in center cell height and the increase in invagination depth, suggesting that this process is closely associated with the redistribution of myosin localization from the apical to lateral regions as invagination progresses.

**Figure 3.**
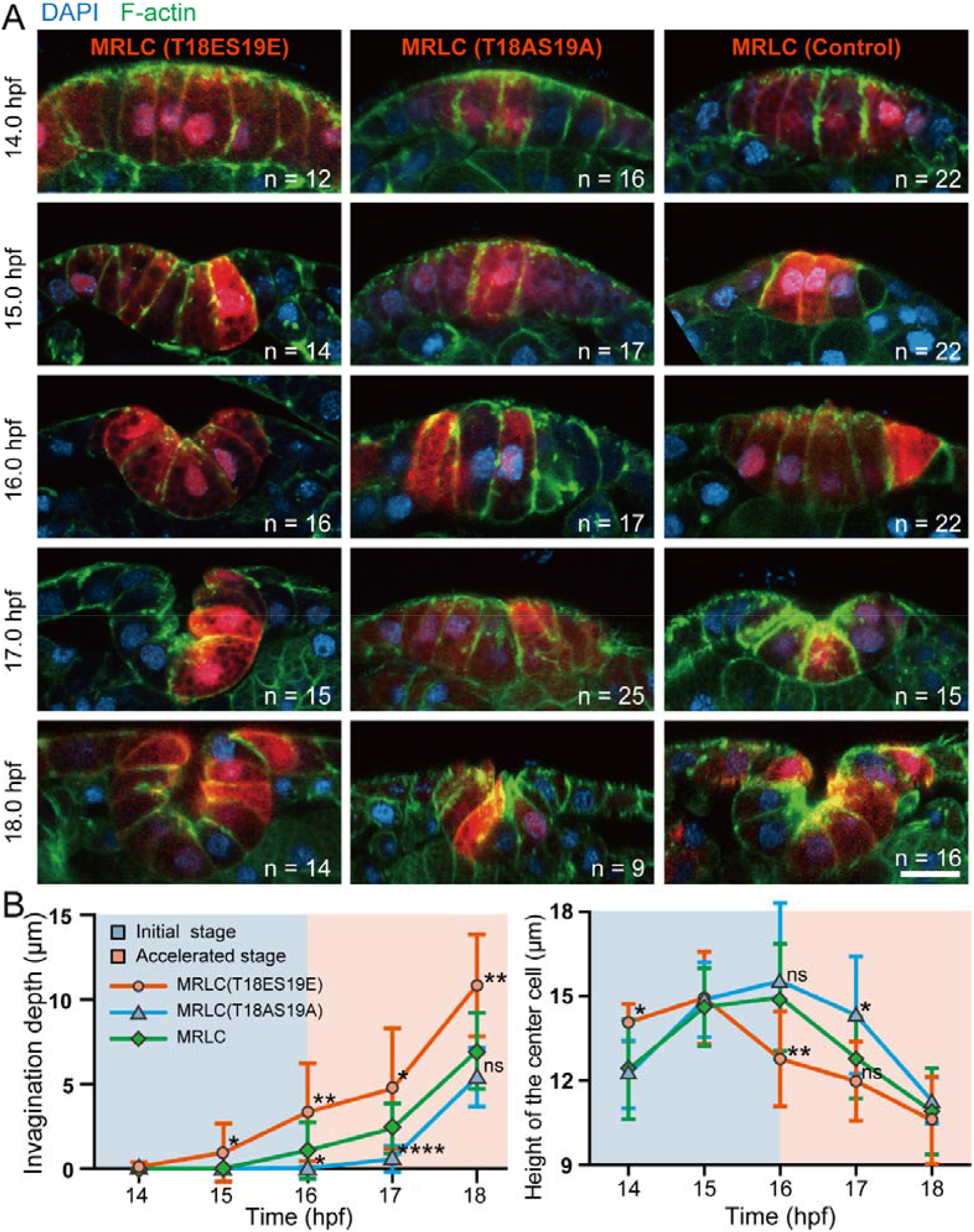
Modulation of atrial siphon invagination by overexpression of myosin mutants. (A) Representative images of *Ciona* atrial siphon primordium expressing wild-type and mutant MRLC constructs. Scale bar: 10 μm. (B) Quantification of invagination depth and the center cell height in MRLC (T18ES19E), MRLC (T18AS19A), and MRLC groups. The blue-shaded region represents the initial stage, while the orange-shaded region indicates the accelerated stage. *p < 0.05, **p < 0.01, ***p < 0.001, ****p < 0.0001.

### Inhibition of myosin activity during apical-to-lateral redistribution impedes invagination progression

To further examine the necessity of the redistribution of myosin between the apical and lateral regions during invagination, we utilized an optogenetic MLCP-BcLOV4(Berlew et al., 2021; Berlew et al., 2020; Berlew et al., 2022; Glantz et al., 2019; Glantz et al., 2018; Yamamoto et al., 2021) system (Figure 4A) to specifically inhibit myosin II activity at the critical redistribution time point between the initial invagination and the accelerated invagination stage (16-17 hpf). This system has been proven to effectively reduce the contractility of epidermal cells in *Ciona* embryos(Qiao et al., 2023). Compared to the control group (Figure 4—figure supplement 1) and dark treatment group, the experimental group with 1 h light exposure, which activated the optogenetic system, blocked further invagination (Figure 4B, Figure 4—video 1, 2), and while invagination ceased to deepen, the center cell height did not show a significant decrease (Figure 4C, D). Immunostaining of active myosin (p-MLC) revealed that after light exposure, lateral myosin intensity was significantly reduced compared to the dark control group, whereas apical myosin levels decreased similarly in both groups (Figure 4—figure supplement 2). This indicates that the optogenetic manipulation effectively attenuates lateral contractility during the accelerated invagination stage, while apical contractility undergoes its normal developmental downregulation (Figure 2B) and shows no significant difference between Light and Dark groups (Figure 4—figure supplement 2B). Together, these results confirm that the redistribution of myosin contractility from the apical to lateral regions, specifically the acquisition of lateral contractility, is essential for the progression of invagination.

**Figure 4.**
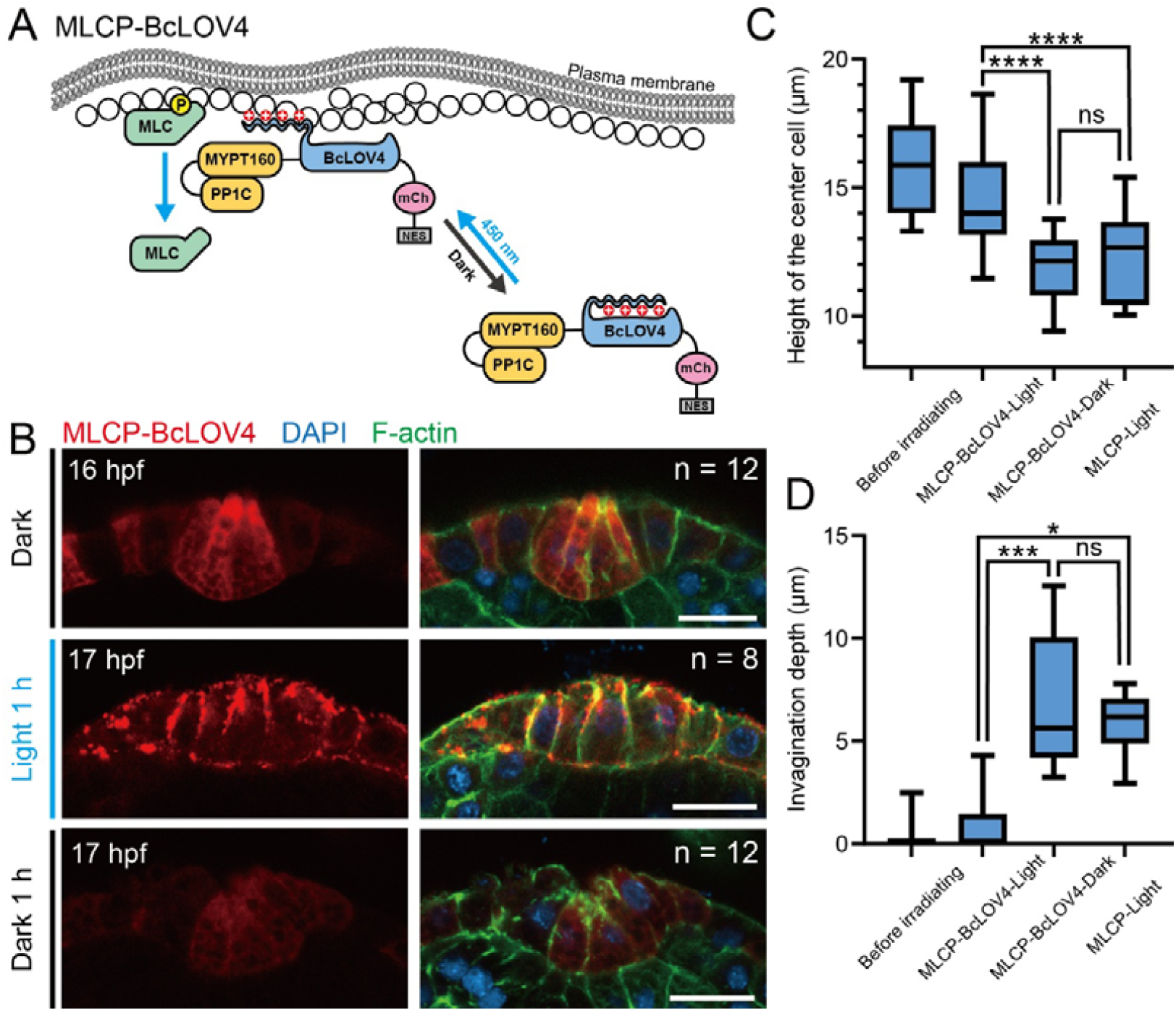
Disruption of contractile forces during rapid invagination of the *Ciona* atrial siphon using an optogenetic system. (A) Schematic diagram depicting the structure and mechanism of the MLCP-BcLOV4 system. The PP1C::MYPT169::BcLOV4::mCherry::NES fusion protein is initially dispersed in the cytoplasm. Upon exposure to blue light, BcLOV4 undergoes a conformational change, allowing it to interact electrostatically with the plasma membrane. This leads to the recruitment of the PP1C and MYPT169 components, subunits of myosin light chain phosphatase (MLCP), to the membrane, where they reduce myosin activity. (B) Representative images of developmental progression in the MLCP-BcLOV4 expression group exposed to blue light for 1_h, and in the dark control group maintained in darkness for 1 h. Scale bar: 10 μm. (C, D) Quantification of invagination depth and the center cell height in the MLCP-BcLOV4 expression group, the dark control group and MLCP control group (Figure 4—figure supplement 1). *p < 0.05, ***p < 0.001, ****p < 0.0001.

### Vertex model simulations recapitulate the mechanical process of ascidian siphon tube invagination

The morphological invagination of epithelial tissues is a conserved mechanical process across diverse developmental contexts. In the early ascidian embryo, a small-scale endoderm plate of only ten cells undergoes invagination via a two-step program of apical constriction and basolateral cell shortening(Sherrard et al., 2010). While the redistribution of actomyosin in this system closely mirrors what we observe in the siphon primordium, their gastrulation-stage movement represents a global evolution of embryonic geometry. In contrast, the invagination of the atrial siphon is a distinctly localized active process within a much larger tissue. A more analogous and extensively studied system is the *Drosophila* ventral furrow formation (VFF), where apical constriction initiates the initial tissue bending(Brodland et al., 2010; Polyakov et al., 2014). The progression into a deep fold further requires lateral contractility(Conte et al., 2012), and is significantly facilitated by in-plane compressive stresses generated by the surrounding tissues(Guo et al., 2022). These studies have established and refined a vertex model framework, effectively elucidating the mechanical underpinnings of such morphogenetic events.

Notably, these classic vertex models of VFF typically incorporate the vitelline membrane—a rigid, non-adhesive extraembryonic shell that tightly encloses the embryo—as a boundary condition(Brodland et al., 2010; Conte et al., 2012; Guo et al., 2022; Polyakov et al., 2014). This membrane acts as a critical external constraint that prevents any outward expansion of the cells. Under such confinement, the mechanical strain generated by active contractility has no freedom degree for outward displacement and can only be released through inward invagination into the yolk. However, in the case of the ascidian siphon, the larval head is enclosed only by a soft, elastic tunic, without such rigid mechanical constraint. In this relatively free-moving tissue, the mechanical mechanism by which a spatiotemporally regulated actomyosin redistribution drives invagination remains to be elucidated.

To evaluate how the bidirectional redistribution of actomyosin drives invagination, we developed a cell-based two-dimensional (2D) vertex model based on previous studies(Guo et al., 2022; Polyakov et al., 2014), and performed simulations for atrial siphon morphogenesis. The 2D cross-section of the *Ciona* embryo head along the transverse plane of its anterior-posterior axis was represented by polygonal cells connected in a circular arrangement, surrounding the inner cells (Figure 5A, Figure 5—figure supplement 1). Since the actomyosin redistribution is highly localized within the surface epithelium, we defined this domain as a local active region occupying an initial *θ*-degree arc of the entire circular tissue. To maintain structural integrity while focusing on epithelial mechanics, the internal cells were simplified as volume-incompressible bulk, providing mechanical support during the invagination process. Cell deformation and tissue morphology were simulated through vertex dynamics, governed by an effective energy functional that accounts for cell area conservation, passive cortical contractility, epithelial bending stiffness, and particularly spatiotemporally regulated active line tensions (Figure 5A, see Methods for details). The redistribution of actomyosin from apical to lateral domains was described by the temporal evolution of normalized intensities 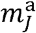 and 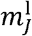 on the apical and lateral edges of cell *J*. Following the normalization of F-actin intensity to the basal surface in experimental measurements (Figure 1—Figure supplement 1A), the model adopted the basal actomyosin intensity 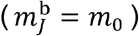 as a non-contractile baseline. The active line tension on the apical or lateral domains was then formulated as 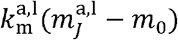, representing the force generated by actomyosin enrichment above the basal baseline, with *k*_m_ serving as the tension coefficient. The intensity *m* within the active region was prescribed by a Gaussian distribution that is centered at the center cell (with index 0) and spans the seven active cells (from index −3 to +3) (Figure 5A, see Methods for details). At 13 hpf,the epithelial cross-section of the larval head comprises approximately 50 cells. Numerical simulations were therefore initiated using a circular epithelium of *N* = 50 cells to examine the morphological processes driven by active tensions (see Methods). Although the tissue expands and its geometry becomes flatter as development proceeds as observed experimentally (Figure 5—figure supplement 1), the size of the active region is much smaller than the total tissue size. Thus, the model focused on the localized impact of actomyosin-driven deformation while omitting global tissue growth. The mechanical implications of tissue size evolution will be discussed further below.

**Figure 5.**
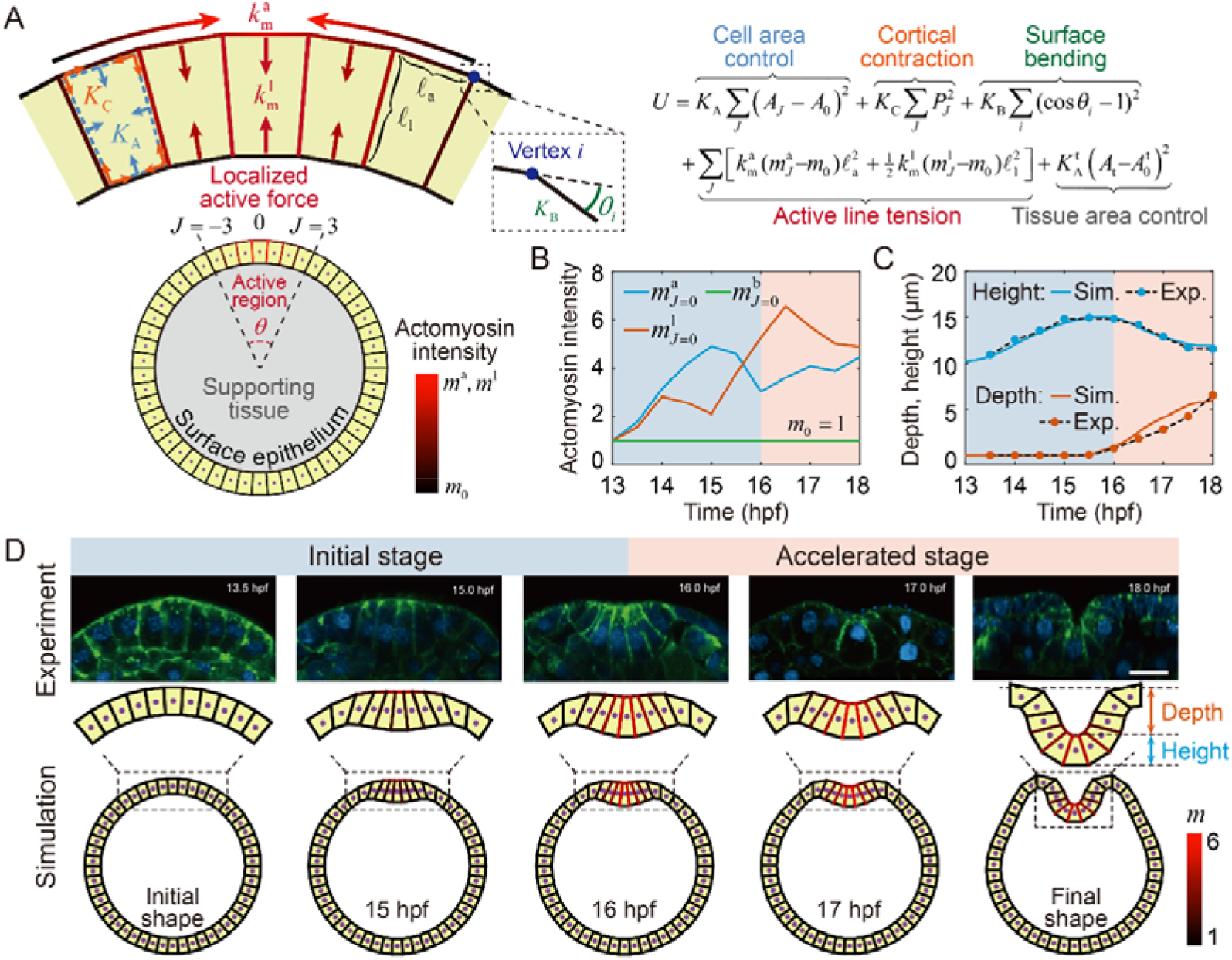
Vertex model simulations of siphon invagination. (A) Schematic of cell-based vertex model. The epithelial tissue is represented by polygonal cells connected in a circular arrangement, surrounding inner supporting cells. The tissue is partitioned into an active region (consisting of cells with indices −3 to +3) with localized active forces, and an inactive region without actomyosin intensity. The effective energy takes into cell area constraint (modulus *K*_A_), passive cortical contraction (coefficient *K*_C_), tissue surface bending (stiffness *K*_B_), tissue area constraint (modulus 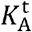), and active line tensions on apical and lateral edges (coefficient *k*_m_). (B) Temporal evolution of actomyosin intensities at apical, lateral, and basal domains of the center cell in the model, parameterized based on experimentally measured F-actin distribution. (C) Evolution curves of invagination depth (orange) and center cell height (blue) in simulations. Dots represent corresponding experimentally measured values. (D) Snapshots at specific timepoints during simulation progression (bottom), compared with corresponding experimental images (up). Colors on cell edges represent the normalized actomyosin intensity.

Our mechanical model successfully reproduced the cell shape evolution and invagination of the atrial siphon observed in experiments (Figure 5C, D, Figure 5 — video 1). During the initial stage, the rapid accumulation of actomyosin on the apical surface in the active region gives rise to apical tension. This localized contraction induces apical narrowing and leads adjacent cells to incline toward the center. The resulting cell elongation breaks the initial apicobasal symmetry, thereby initializing the invagination into the basal side during 15-16 hpf. After 16 hpf, the lateral actomyosin intensity increases significantly and induces high contractility on the lateral surface, pulling cells and shortening the cell height. The synergy between apical and lateral contraction further drives the invagination process, ultimately establishing the characteristic siphon-like epithelial architecture (Figure 5D). The invagination depth is quantified as the vertical distance from the apical surface of the center cell to a baseline connecting the apical junctions of cells ±4 and ±5, and the height of the center cell is defined as the vertical distance between its apical and basal surfaces. Both quantifications obtained from the simulations are consistent with the experimentally measured data (Figure 5C), which confirms the validity of our model in capturing the mechanical process of ascidian siphon tube invagination.

Note that, we find that “puckering” shape occurs surrounding the invagination in simulations (Figure 5D). Previous vertex models of VFF simulating cells within a rigid unmovable boundary do not exhibit this effect, and the invaginating cells remain tightly adherent, collapsing into a solid structure rather than forming a hollow tube(Conte et al., 2012; Polyakov et al., 2014). However, in our system of siphon morphogenesis, a tubular structure ultimately forms in the epithelium without strong boundary constraints. The final tubular structure is accompanied by significant F-actin accumulation at the apical surface (Figure 1A), which provides mechanical support and bending resistance(Hirashima & Matsuda, 2024; Wyatt et al., 2020). Thus, this “puckering” in our simulations arises from the bending energy, which enforces smooth curvature transitions along the circular surface. Actually, similar mild puckering is also observed *in vivo* at early stages (∼17 hpf in Figure 1A), when the tissue size is small, but it rapidly disappears as the tissue grows and the geometry becomes flatter. Therefore, in our model, this shape discrepancy is expected to diminish as the mean curvature decreases in a larger system (see Figure 6D). In addition, incorporating the cell-cell adhesion between the surface epithelium and internal bulk cells likely further suppresses this puckering in vivo, as outward evagination would require the coordinated deformation of underlying tissues.

**Figure 6.**
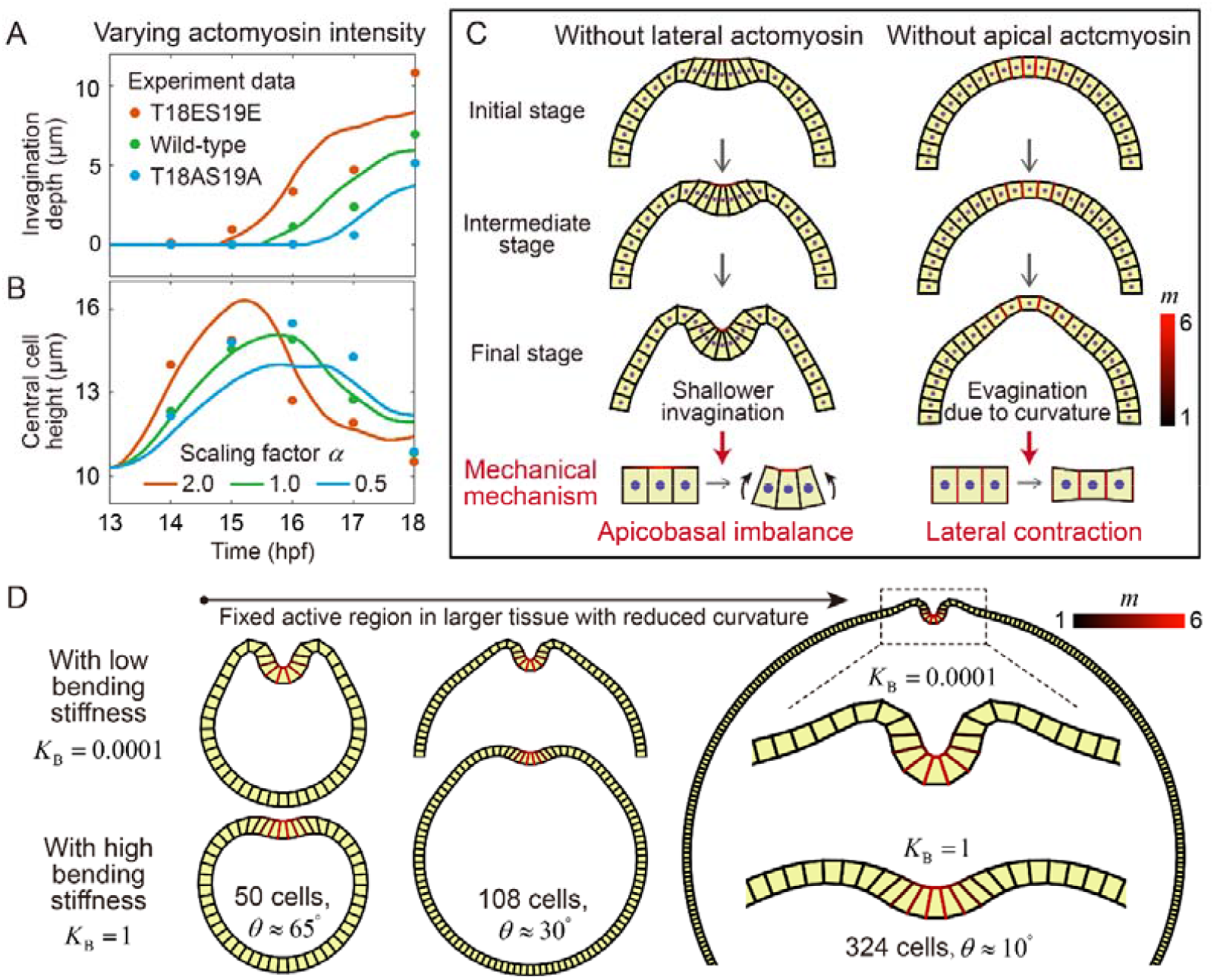
Mechanics of siphon invagination revealed by vertex model simulations. (A, B) Invagination depth (A) and center cell height (B) regulated by changes in actomyosin intensities under the scaling factor *α* at both apical and lateral regions of all active cells. The data points are from the mutant experiments in Figure 3B. (C) Contributions of apical and lateral contractility. Simulations lacking lateral (left) or apical (right) activity result in shallower invagination or outward evagination, respectively, which highlight the distinct roles of apical constriction in initiating apico-basal imbalance and lateral contraction in driving deep folding. (D) Effects of tissue size and bending stiffness. With the active region size held constant, the global tissue curvature decreases as the total cell number increases. For *N* = 108, only the upper half of the tissue at *K*_B_ = 0.0001 is shown. For *N* = 324, a portion of the tissue is shown at *K*_B_ = 0.0001, and only the region near the active region is shown at *K*_B_ = 1.

Furthermore, we varied the apico-lateral actomyosin intensities *m* = *am*_exp_ by introducing a scaling factor *α* to the experimental measured value *m*_exp_ into the model, to simulate the unphosphorylated or diphosphorylated mutants with altered myosin activity (Figure 3). As expected, enhancement of both apical and lateral actomyosin activities increased the invagination depth while reducing the height of the center cell, and vice versa (Figure 6A,B). These trends are consistent with the experimental mutant data (Figure 3B). That being said, the simulated invagination depth remains smaller than that observed experimentally across all conditions, and the invagination speed (represented by slopes of the curves in Figure 5C) gradually slows down at later stages, in contrast to experiments. We attribute this discrepancy to the absence of growth-induced compressive stress in the model: as the invagination initiates, the furrow may act as a release site for accumulated internal stress, thereby accelerating its progression. This hypothesis requires validation through quantitative characterization of tissue size and stress distribution in the future. In addition, the final cell height in simulations is slightly larger than measured experimentally, because the model does not account for the mechanical constraints imposed by the growth of internal supporting tissues. Despite these differences, the results demonstrate that actomyosin contractility controls the magnitude of active forces and thereby regulates the morphology of invagination.

### Coupled mechanism of apicobasal tension imbalance and lateral contraction underlying epithelial invagination

To theoretically distinguish the effects of apical and lateral contraction on morphogenesis, we next performed simulations in which the two processes are considered independently. We considered active forces arising from either apical or lateral contraction alone, while maintaining the same actomyosin intensity profiles independently as in Figure 5B. We found that the apical contractility alone is sufficient to induce invagination, but produces a shallower invagination and taller cell shape in the absence of lateral actomyosin (Figure 6C). In contrast, without apical actomyosin, lateral contractility alone shortens the cells but cannot reproduce invagination. In a curved tissue with a surface bending modulus, such localized height reduction is more easily accommodated by bending outward. As a result, instead of driving invagination, lateral contraction biases the tissue toward outward deformation, leading to evagination (Figure 6C). These results suggest two mechanical mechanisms underlying the siphon invagination: an effective bending moment generated by apico-basal tension imbalance drives tissue symmetry breaking and determines the location and direction of buckling, whereas lateral actomyosin-driven contraction regulates cell shape and facilitates tissue invagination.

After separately identifying the mechanical roles of apical and lateral contractility, we further examined the mechanical consequence of actomyosin redistribution during morphogenesis. We modified the temporal actomyosin dynamics by maintaining apical activity during the accelerated stage or prematurely enhancing lateral activity during the initial stage. When apical actomyosin activity was maintained after 16 hpf instead of decreasing during the accelerated stage (Fig. 6—figure supplement 1A), the early deformation process before 16 hpf remained unchanged. However, sustained apical tension induced stronger cell elongation, while the final invagination depth remained comparable to the control simulation (Fig. 6—figure supplement 1B,C). In contrast, elevating lateral actomyosin activity during the initial stage (14-16 hpf) suppressed central cell elongation and the inward movement of surrounding cells toward the central cell, resulting in earlier bending deformation with a flatter invaginating morphology (Fig. 6—figure supplement 1D-F). Although both perturbations eventually converged to similar final shapes, their distinct morphogenetic trajectories demonstrate the mechanical importance of the actomyosin redistribution sequence. The initial dominance of apical contractility, with limited lateral contraction, first promotes cell elongation and convergence of the active region, whereas the subsequent shift toward lateral contractility enables efficient cell shortening and deep tissue invagination.

Finally, we investigate the proposal local active mechanisms under varying tissue curvature and bending stiffness. In the initial configuration (*N* = 50 cells), the active region spans approximately *θ* ≈ 65°. However, in developing tissues, the length scale of the whole tissue is much larger than that of the active region. To examine this, we fixed the size of the active region (seven cells) and constructed larger systems with *N* = 108 (*θ* ≈ 30^°^) and *N* = 324 (*θ* ≈ 10^°^), while keeping all other parameters unchanged. The resulting final tissue shapes are shown in Figure 6D. At low bending stiffness (*K*_B_ = 0.0001), pronounced local invagination emerges robustly across tissues with different curvature and system sizes, indicating that siphon invagination is governed by a highly localized active process that requires only apico-lateral contractions in a small group of cells. Notably, in the larger system with 324 cells, the puckering surrounding the invagination is significantly reduced because bending constraints are more prominent in highly curved tissues. As the tissue becomes flatter at later developmental stages, such puckering is therefore not expected to be observed experimentally. When the bending stiffness is increased (*K*_B_ = 1), local active contraction induces only cell shape changes and fails to drive invagination. This reflects a competition between local active contractile forces and the global mechanical resistance of tissue, suggesting that sufficiently strong active forces are required to overcome bending constraints and reshape tissue morphology. We note that the present model does not exclude the potential contribution of growth-induced compressive stresses, which may arise from tissue growth and augment the active mechanisms described above to accelerate invagination at later stages. This interplay will be further elucidated in future work.

## Discussion

The morphogenesis of the *Ciona* atrial siphon provides a compelling model to dissect how biomechanical forces orchestrate tissue invagination. Our study revealed that actomyosin contractility underwent a dynamic spatial redistribution during this process, initially redistributing from the lateral cortex to the apical region to promote apical constriction and establish the initial cell shape, and subsequently redistributing back to the lateral regions to promote center cell shortening and accelerate the tissue invagination. By combining actomyosin localization analysis, genetic perturbations, optogenetic manipulation, and vertex model, we established a mechanistic framework linking force dynamics to tissue remodeling. This work not only advances our understanding of siphon morphogenesis but also offers broader insights into the principles governing force-driven tissue shaping in developing organisms.

Sequential activation of actomyosin contractility has been documented in several epithelial folding systems, yet our study reveals distinct features. In ascidian endoderm invagination, actomyosin shifts from apical to basolateral regions(Sherrard et al., 2010), whereas our system exhibits bidirectional redistribution between apical and lateral domains, with the basal domain playing a passive role. More notably, during *Drosophila* ventral furrow invagination, lateral contractility is not essential for the second folding phase(Guo et al., 2022); in contrast, our optogenetic inhibition demonstrates that lateral contractility is obligatory for the accelerated stage. These comparisons establish bidirectional actomyosin redistribution as a distinct mechanical paradigm for sequential morphogenesis.

Our findings demonstrated that the atrial siphon invagination was driven by a coupled mechanism including apicobasal tension imbalance and lateral contraction in a two-stage process. During the initial stage (13.5-16.0 hpf), actomyosin redistributed from the lateral cortex to the apical cortex, generated contractile forces that induced apicobasal tension imbalance and reduced apical cell area and elongated the primordium. A transient actomyosin ring surrounding the primordium is prominent at the early initial stage and may cooperate with apical constriction, but it diminishes during the accelerated stage, indicating that sustained compression is not required. This aligns with classical models of epithelial folding, where apical actomyosin networks drive tissue curvature through localized contraction, often accompanied by an increase in cell height, as observed in the invagination of the *Drosophila* embryonic trachea(Schottenfeld et al., 2010) and salivary gland placode(Pearl et al., 2017). Future experiments using laser ablation or optogenetic inhibition specifically targeting this actomyosin ring could help dissect its precise contribution during the early invagination stage. Additionally, in traditional invagination models, various epithelial tissues use different mechanisms to drive deeper invagination. For example, in *Drosophila* tracheal development, mitosis acts as a critical turning point in accelerating invagination(Kondo & Hayashi, 2013). During the early slow phase of tracheal invagination, apical constriction under EGFR signaling forms a shallow pit. As invagination transitions into a rapid phase, mitotic entry induces cell rounding, which increases tension and epithelial buckling, thereby accelerating the invagination process and facilitating the internalization of placode cells(Kondo & Hayashi, 2013; Nishimura et al., 2007). Moreover, cell apoptosis can also expedite the invagination process, as seen in *Drosophila* leg morphogenesis, where actomyosin cables formed in apoptotic cells help trigger the invagination(Kiehart, 2015; Manjón et al., 2007; Monier et al., 2015). In Drosophila ventral furrow formation, in-plane compressive stresses generated by surrounding tissues significantly facilitate the deep fold(Guo et al., 2022). Distinct from these classical models, the accelerated invagination stage of *Ciona* atrial siphon morphogenesis was driven by a critical redistribution of apical actomyosin to the lateral regions, providing a force-generating mechanism for invagination progression without relying on cell proliferation or apoptosis to accelerate the process. The lateral enrichment of p-MLC correlated with a reduction in center cell height and the initiation of invagination, suggesting that lateral forces actively pulled the tissue inward. This spatial redistribution of contractile forces in *Ciona* atrial siphon invagination is reminiscent of mechanisms observed in certain epithelial folding processes, such as the *Drosophila* wing disc, where invagination is not solely driven by apical constriction but can also result from increased lateral tension or reduced basal tension(Sui et al., 2018). Our optogenetic inhibition of myosin activity during the apical-to-lateral redistribution—using the MLCP-BcLOV4 system—directly confirmed that this redistribution was not merely a passive consequence of invagination but was an active driver of the process.

Our study revealed a strong coupling between cellular shape changes and tissue remodeling during *Ciona* atrial siphon invagination, where the reduction in center cell height was closely associated with invagination depth. Tissue morphogenesis emerges from the integrated actions of multiple cellular behaviors, such as contraction, adhesion, and migration, that work in concert to remodel epithelial architecture(Chan et al., 2017). The transmission of local cell-generated forces across tissues is further shaped by mechanical cues and geometric constraints, enabling the transformation of cellular-scale changes into organ-scale morphogenesis(Collinet & Lecuit, 2021). By perturbing myosin activity, we found that this coupling was directly regulated: dominant-negative myosin mutants delayed both processes, whereas hyperactive myosin accelerated invagination. The quantitative agreement between vertex model analysis and experimental measurements validated the critical role of actomyosin contractility dynamics in linking cellular shape changes to tissue remodeling. Furthermore, the model also uncoupled the effects of the actomyosin in different domains, and predicted that stronger apical-lateral redistribution facilitates the rapid reduction in height of invaginating cell. The *Ciona* atrial siphon, which underwent invagination without cell proliferation or apoptosis, provided a minimal system for investigating the role of biomechanical forces in tissue remodeling. Our findings emphasized that the deformation of a few key cells under spatially precise mechanical regulation, along with their coordination with surrounding tissues, is crucial for proper morphogenesis.

While this study clarified the role of actomyosin redistribution in atrial siphon invagination, several open questions remain regarding the molecular mechanisms underlying this process. One key question is what molecular signals drive the apical-to-lateral redistribution of contractility. Possible candidates include Rho GTPase pathways(Martin et al., 2016), which regulate myosin activation; apical-basal polarity(Peng et al., 2020), which coordinate actin reorganization during morphogenesis; and mechanosensitive ion channels that respond to tissue strain(Jin et al., 2020). Additionally, the role of cell-cell adhesion and extracellular matrix remodeling in mediating force transmission during invagination requires further investigation. Using tension sensors or perturbing adhesion molecules may provide new insights into these interactions. In addition, our experimental system involves significant global tissue remodeling: as development proceeds, the number of cells increases and the tissue expands substantially, accompanied by classic growth-induced compressive stress(Guo et al., 2022). Although we did not directly measure such stress in the siphon primordium, we observed a progressive flattening of the overall tissue geometry, which likely facilitates invagination and lumen formation. While our experiments and vertex model confirm that active tension alone is sufficient to drive invagination, tissue growth and the associated mechanical constraints may play critical roles in achieving the final tubular architecture. Future studies should integrate tissue growth and mechanosensitive feedback into theoretical models, and experimentally quantify how growth-induced stresses cooperate with local actomyosin redistribution to robustly shape the atrial siphon. Although our vertex model effectively clarified the contribution of force redistribution to invagination, it does not account for tissue growth or the potential for mechanosensitive feedback on the generation and redistribution of actomyosin. Future theoretical developments should focus on integrating tissue growth and mechanochemical feedback linking molecular signaling with tissue-level deformation to explain robust morphogenesis.

## Materials and Methods

### Experimental animals’ preparation and electroporation

*Ciona* adults were collected from the coast of Qingdao and Rongcheng, Shandong, China. They were maintained in the laboratory seawater circulation system under constant light conditions to maintain their stability. Mature eggs and sperm were separately collected from the dissected adults and subsequently mixed for fertilization. The Fertilized eggs were dechorionated in seawater containing 1 % sodium thioglycolate (T0632; Sigma), 0.05 % protease E (P5147; Sigma) and 0.032 M NaOH. The dechorionated eggs were then used for plasmid electroporation based on the previous technique procedure(Christiaen et al., 2009). Finally, the embryos were cultured in an agar-coated dish with microporous-filtered seawater in 18□ for further observation. Detailed key resource information is annotated in Appendix 1—table 1.

### Plasmid construction

*Ciona* myosin II light chain (MRLC) was amplified with primers MRLC-F and MRLC-R (Table S1). MRLC-mCherry(Dong et al., 2011); MRLC(T18AS19A)-mCherry and MRLC(T18ES19E)-mCherry(Denker et al., 2015); MLCP-BcLOV4 and MLCP(Qiao et al., 2023) were amplified with the primers listed in Appendix 1—table 2. For embryos expressing MRLC constructs, only those in which the center cell and more than half of the surrounding cells in the primordium showed clear mCherry fluorescence were selected for quantification.

### Immunofluorescence

*Ciona* embryos were fixed with stationary liquid(Sherrard et al., 2010), which consisted of 100 mM HEPES (pH 6.9); 100 mM EGTA (pH 7.0); 10 mM MgSO4; 2% formaldehyde; 0.1% glutaraldehyde; 300 mM dextrose and 0.2% Triton X-100 for 40 m at room temperature. Following fixation, the embryos were washed three times with PBS and subsequently incubated in PBST (PBS supplemented with 0.1% Triton X-100) for 30 m to enhance permeability. To reduce autofluorescence, embryos were treated with 0.1% sodium borohydride in PBS for 20 m at room temperature. For immunostaining, embryos were incubated with a 1:250 dilution of Phospho-Myosin Light Chain 2 (Ser19) antibody (#3671: Cell Signaling) at room temperature for 24 h. After three additional washes with PBS, a 1:200 dilution of Alexa Fluor 568 anti-Rabbit IgG (A11011: Invitrogen) was added and incubated at room temperature for 48 h. For cell boundary visualization, embryos were stained with Alexa Fluor 488 Phalloidin (A12379: Invitrogen) at a 1:200 dilution. Finally, after three washes with PBS, embryos were mounted in DAPI-containing mounting medium and prepared for imaging.

### Imaging and optogenetics

Live imaging, photoactivation experiments, and image acquisition were performed using a Zeiss LSM 980 confocal microscope (Carl Zeiss). For optogenetic experiments, *Ciona* embryos were placed in a 35 mm glass-bottom dish for imaging. To activate the optogenetic system, the designated region of interest was exposed to a 488 nm laser for 1 h. The control group for dark treatment was placed in a dark box at the same temperature for 1 h.

### Quantification and statistical analysis

All images were analyzed and quantified using ImageJ (version 1.54p, NIH) and Imaris (version 9.0.1, Bitplane). For intensity measurements of active myosin (p-MLC) and F-actin, the apical domain was defined as a segmented line (width 1 μm) drawn along the apical cell⍰cell junctions of the center cell; the lateral domain was defined as a line along the lateral membranes excluding the apical and basal regions; the basal domain was defined as a line along the basal surface. Fluorescence intensities were normalized to the basal intensity of the same center cell (set to 1). Statistical analyses were performed using GraphPad Prism 10.1.2. Data are presented as mean ± SEM. Comparisons between two groups were analyzed by two⍰tailed Student’s t⍰test. Significance levels are indicated as *p < 0.05, **p < 0.01, ***p < 0.001, ****p < 0.0001; ns indicates not significant. Sample sizes (n) for each experiment are indicated in the corresponding figure legends.

### Vertex model simulations

#### Model description

The 2D cross-section of the Ciona embryo head along the transverse plane of its anterior-posterior axis is represented by polygonal cells connected in a circle, surrounding the inner supporting tissue (Figure 5A). Each polygon is defined by a set of vertices and edges. Vertices on the apical surface of cells are subject to free boundary conditions, which differs from the previous models(Conte et al., 2012; Polyakov et al., 2014). We define the invagination domain as a local active region occupying an initial θ-degree arc of the entire circular tissue. To maintain structural integrity while focusing on epithelial mechanics, the internal cells are simplified as volume-incompressible bulk, providing mechanical support during the invagination process. Cell deformation and tissue morphology are simulated through vertex dynamics, governed by an effective energy functional

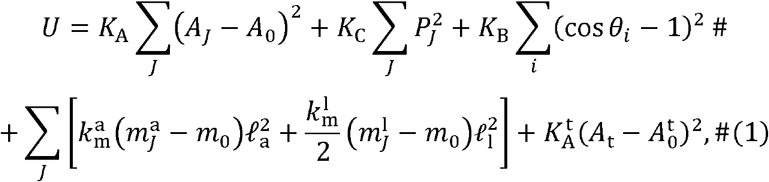

where the five terms result from, respectively, cell area control, cell passive cortex contractility, tissue surface bending energy, active line tensions on apical and lateral surfaces, as well as tissue area constraint. *K*_A_ is the cell area modulus, and *A*_*J*_ and A_0_ are the current and preferred areas of the cell *J*, respectively. 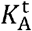 is the tissue area modulus, and *A*_t_ and 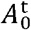 are the current and preferred areas of the supporting tissue. *K*_C_ denotes the passive cortical contraction coefficient of cells and *P*_*J*_ represents the cell perimeter. *K*_B_ denotes the bending stiffness of the cell monolayer, representing its resistance to curvature changes. Physically, this stiffness arises from the recruitment of F-actin and other cytoskeletal elements at the tissue surface, serving as a critical mechanical constraint that ensures structural stability under free boundary conditions. θ_*i*_ is the angle between the neighboring apical or basal edges of vertex *i* (Figure 5A). 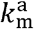 and 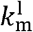 are the tension coefficient of actomyosin (*m*^a^ and *m*^l^) recruited the apical and lateral surfaces whose lengths are *ℓ*_a_ and *ℓ*_l_, respectively. Experimental measurements of F-actin intensity on apical and lateral surfaces were normalized to the basal surface (Figure 1—Figure Supplement 1A). Consequently, we set the basal actomyosin intensity to 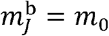, serving as a non-contractile baseline. The active line tension on the apical or lateral domains is then formulated as 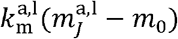, representing the force generated by actomyosin enrichment above this basal baseline. Notably, since each lateral edge is shared by two neighboring cells, the coefficient 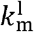 is halved in the force calculation for each individual cell. ∑_*J*_ stands for the summation over all cells and ∑_*i*_ runs over all apical and basal vertices.

Based on experimental measurements, the F-actin signal intensity is highest in the central cell and rapidly attenuates in the neighboring cells. To capture this spatiotemporal evolution within the active region, the actomyosin intensity is prescribed by a Gaussian distribution centered at the center cell (*J* = 0). The intensity for any cell *J* is defined as

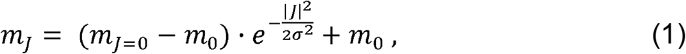

where m_*J* =0_ is the experimental value measured from the central cell (as shown in Figure 5B), and *J* denotes the cell index extending from the center. The parameter *σ* controls the width of the active zone. In accordance with experimental observations, we set *σ* = 1.8. This distribution effectively spans from *J* = −3 to +3, encompassing approximately seven active cells, while the actomyosin activity is set to *m*_0_ for cells outside this region (|*J*| > 3).

#### Simulation scheme

In the simulations, we set the cell number *N*= 50 and a free boundary condition. The central seven cells (index −3 to +3) are considered as active cells under contractile forces whose actomyosin intensity varies in different stages. The other cells are observed to be inactive whose actomyosin intensities are always equal to *m*_0_ . In our simulations, tissue morphogenesis is represented by vertices motion and cell deformation. The motion of vertex *i* obeys the Langevin equation

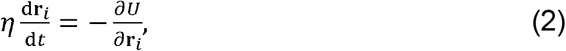

where *η* denotes the friction coefficient. The equation was numerically solved using the forward Euler method with a time step of 0.001, using MATLAB R2021a. Simulations are run until *t* = 500, which corresponds to 18 hpf in experimental development, given a characteristic time scale of *τ* = 36 seconds. The final tissue shape is obtained after the system has relaxed to a steady-state configuration. The length scale *L* = 8.91 µm is determined by the normalization of initial cell height *h*_0_ in model under real center cell height. The other parameters are all normalized with the time scale *τ*, length scale *L*, and friction coefficient *η* . Dimensionless parameters are set as follows: 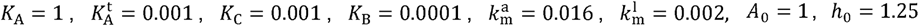, and 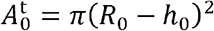, where *R*_0_ = *NA*_0_/(2 *π h*_0_) is the radius of initial tissue configuration. These parameter values are determined by numerical simulations under which the simulated invagination depth and center cell height imitates the experimental observations.

## Acknowledgements

We are grateful to all members of the Fang Zongxi center for helpful discussions.

## Author contributions

B.D., B.L. J.Q., P.Y. conceived of the study; B.D., J.Q., P.Y., B.L., H.P., and W.S. analyzed data; P.Y. and B.L. designed the vertex model; B.D., J.Q., P.Y. and B.L. wrote the original draft. All authors approved the final version of the article.

## Data and resource availability

### Lead contact

Requests for further resources should be directed to, and will be fulfilled by, the lead contact, Bo Dong.

### Materials availability

All unique/stable reagents generated in this study are available from the lead contact with a completed materials transfer agreement.

### Data and code availability

Model code and data are available upon request, and any additional information required to reanalyze the data reported in this paper is available from the lead contact upon request.

## Figure legends

**1—figure supplement 1.**
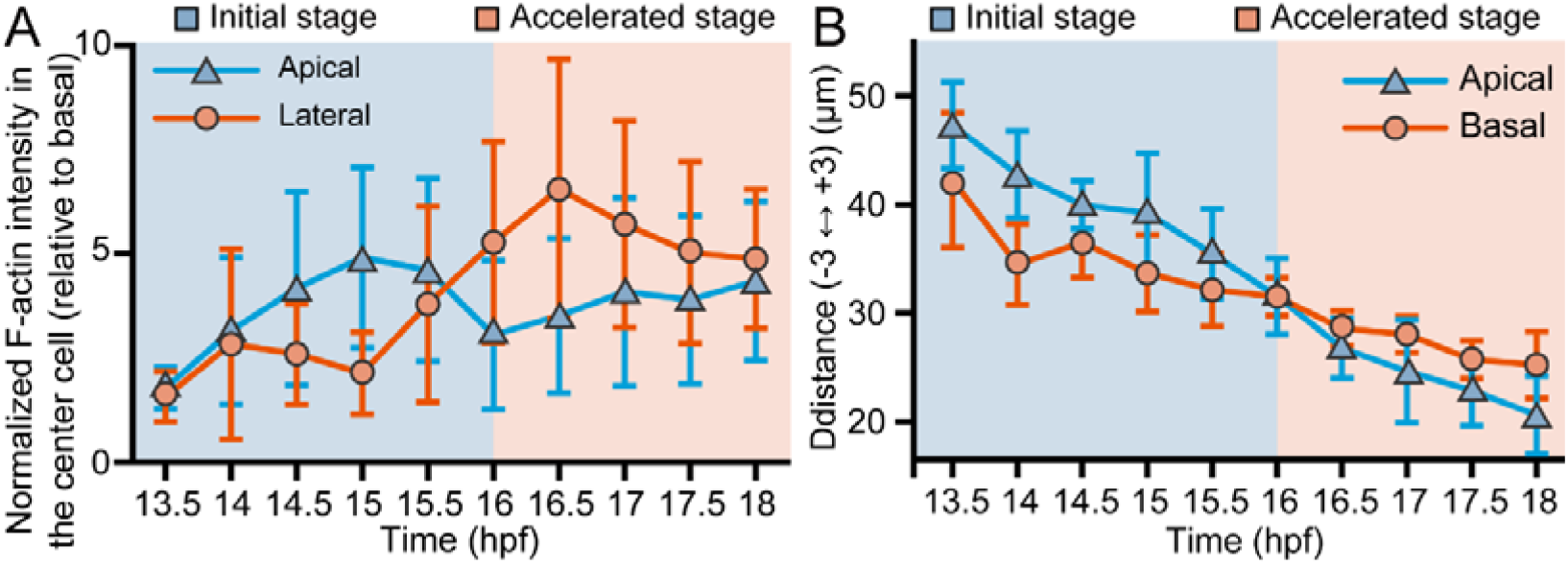
Quantification of F-actin intensity and intercellular distance during *Ciona* atrial siphon morphogenesis. (A) Normalized F-actin intensity at the apical and lateral regions of center cells during *Ciona* atrial siphon morphogenesis (basal level set to 1). (B) Quantification of the linear distance between the -3/-4 and +3/+4 cell junctions at the apical or basal surface in the atrial siphon of *Ciona* embryos. The blue-shaded region represents the initial stage, while the orange-shaded region indicates the accelerated stage. Representative images are shown in Figure 1A. n = 20.

**1—figure supplement 2.**
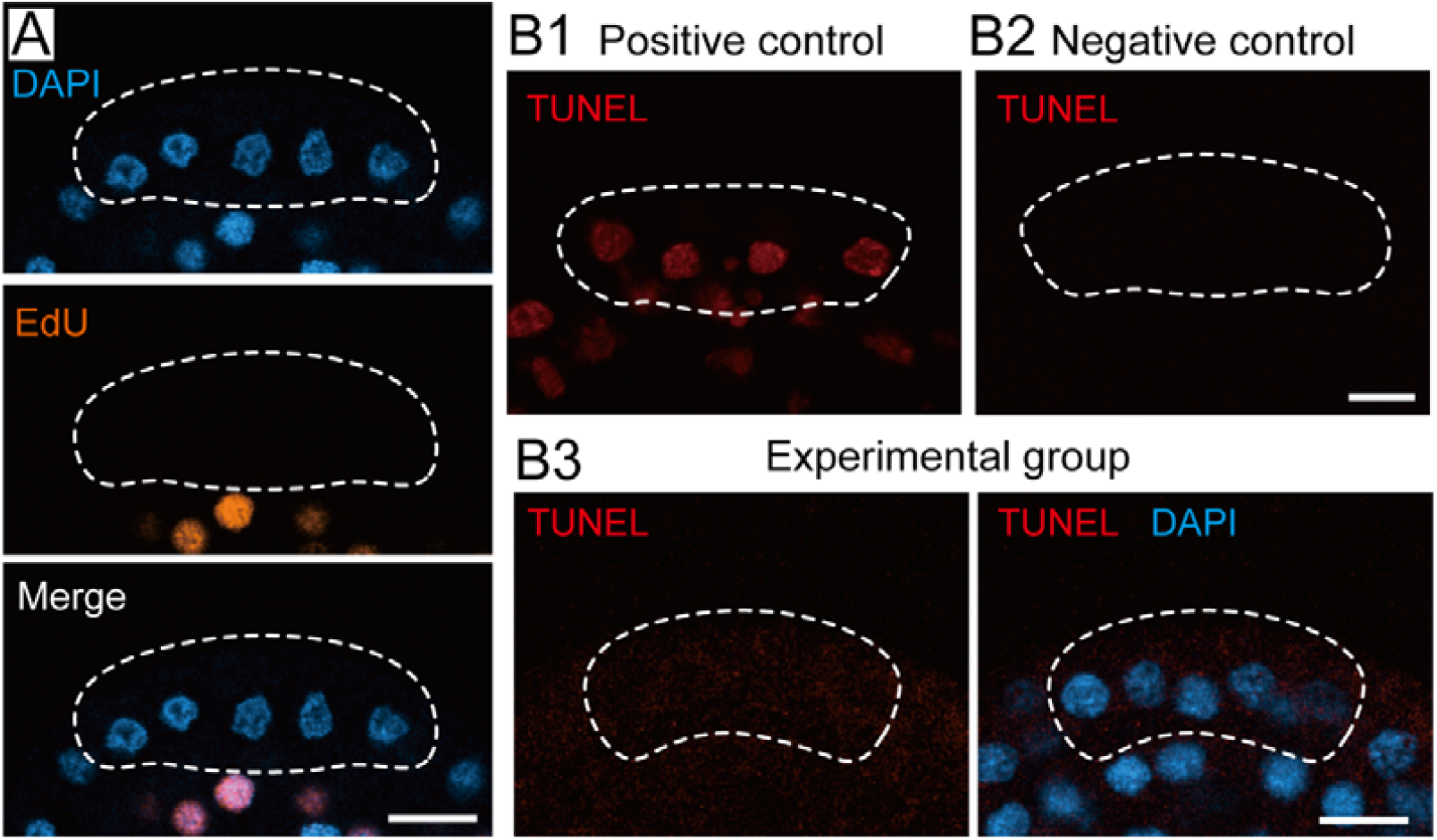
EdU and TUNEL staining during *Ciona* atrial siphon morphogenesis. (A) Representative images of EdU staining at 14-15 hpf. Orange: EdU-positive nuclei indicating cell proliferation. Blue: DAPI. No EdU signal was detected in the atrial siphon primordium (white dashed outline). Scale bar: 10 μm. n = 10. (B) Representative images of TUNEL staining at 15 hpf. (B1) Positive control: DNase I pretreatment (20 U/mL, 10 min) induced DNA fragmentation. (B2) Negative control: staining performed without terminal deoxynucleotidyl transferase (TdT) enzyme. (B3) Experimental group: no detectable TUNEL signal in the atrial siphon primordium (white dashed outline). Scale bar: 10 μm. n = 10.

**4—figure supplement 1.**
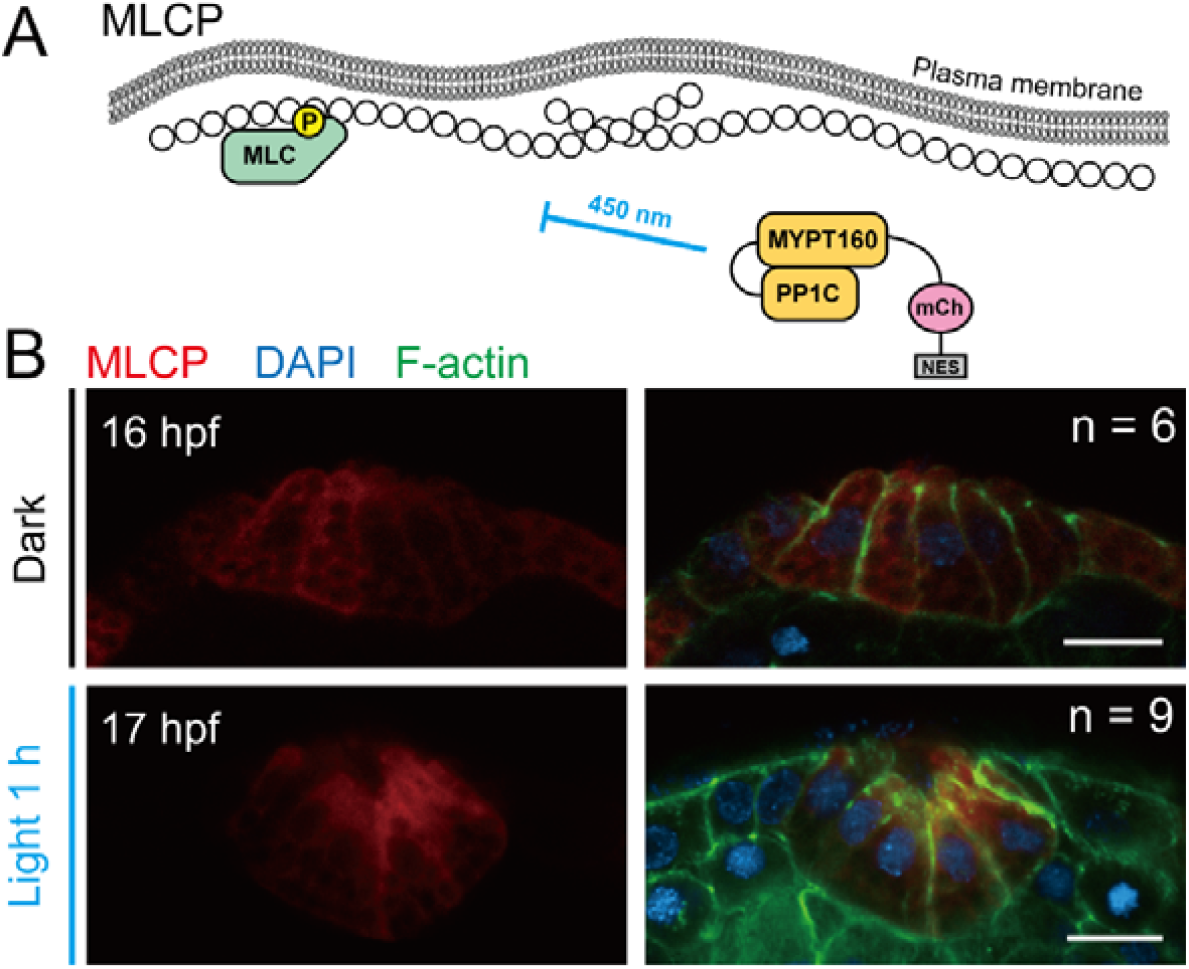
MRCL control group in the optogenetic experiment. (A) Schematic diagram depicting the structure and mechanism of the MLCP control system. The PP1C::MYPT169::mCherry::NES fusion protein remained diffuse in the cytoplasm under light exposure and failed to function. (B) Representative images of developmental progression in the MLCP control group exposed to blue light for 1_h. Scale bar: 10 μm.

**4—figure supplement 2.**
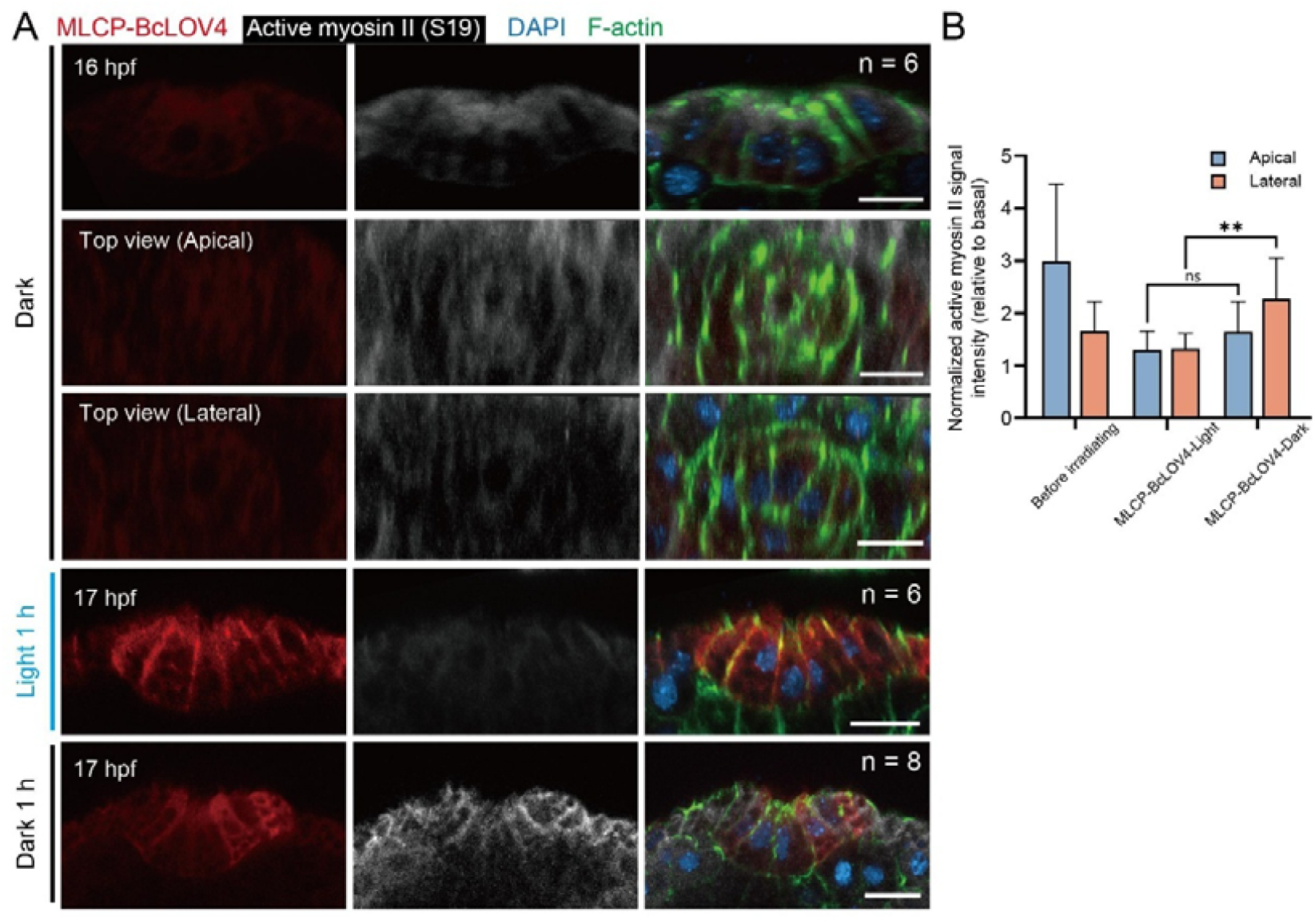
Optogenetic inhibition effectively reduces the activity of lateral myosin. (A) Representative images of active myosin II (anti-pS19 MRLC,red) and F-actin (green) staining in the MLCP-BcLOV4-expressing embryos after 1 h of light exposure or dark control. Scale bar: 10 μm. (B) Quantification of normalized active myosin II intensity at the apical and lateral domains of the center cell. **p < 0.01.

**Figure 4—video 1**. The developmental processes of the MLCP control group under blue light illumination for 1 h. Scale bar: 10 μm.

**Figure 4—video 2**. The developmental processes of MLCP-BcLOV4-expressed group under blue light illumination for 1 h. Scale bar: 10 μm.

**5—figure supplement 1. .**
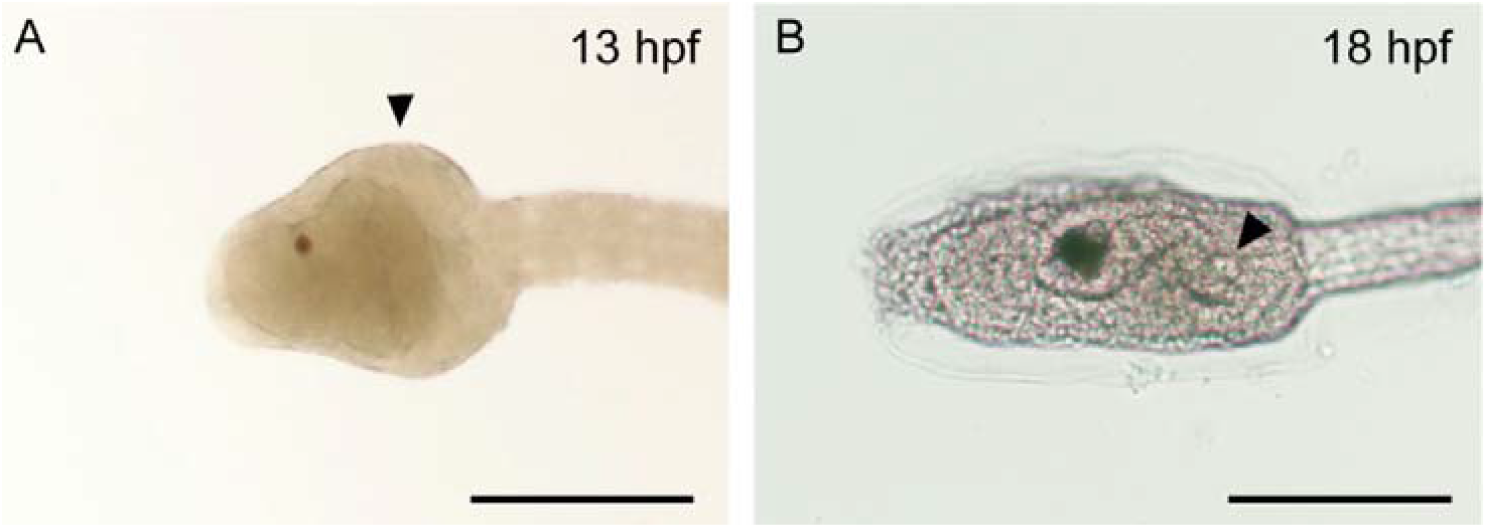
Entire head morphology of *Ciona* embryo. (A, B) Bright⍰field images showing the entire head of *Ciona* embryos at 13 hpf and 18 hpf, illustrating how the tissue geometry becomes flatter during growth. Black arrow indicates the position of atrial siphon. Scale bar: 100 μm.

**Figure 5—video 1**. Vertex model simulation of atrial siphon invagination.

**6—figure supplement 1.**
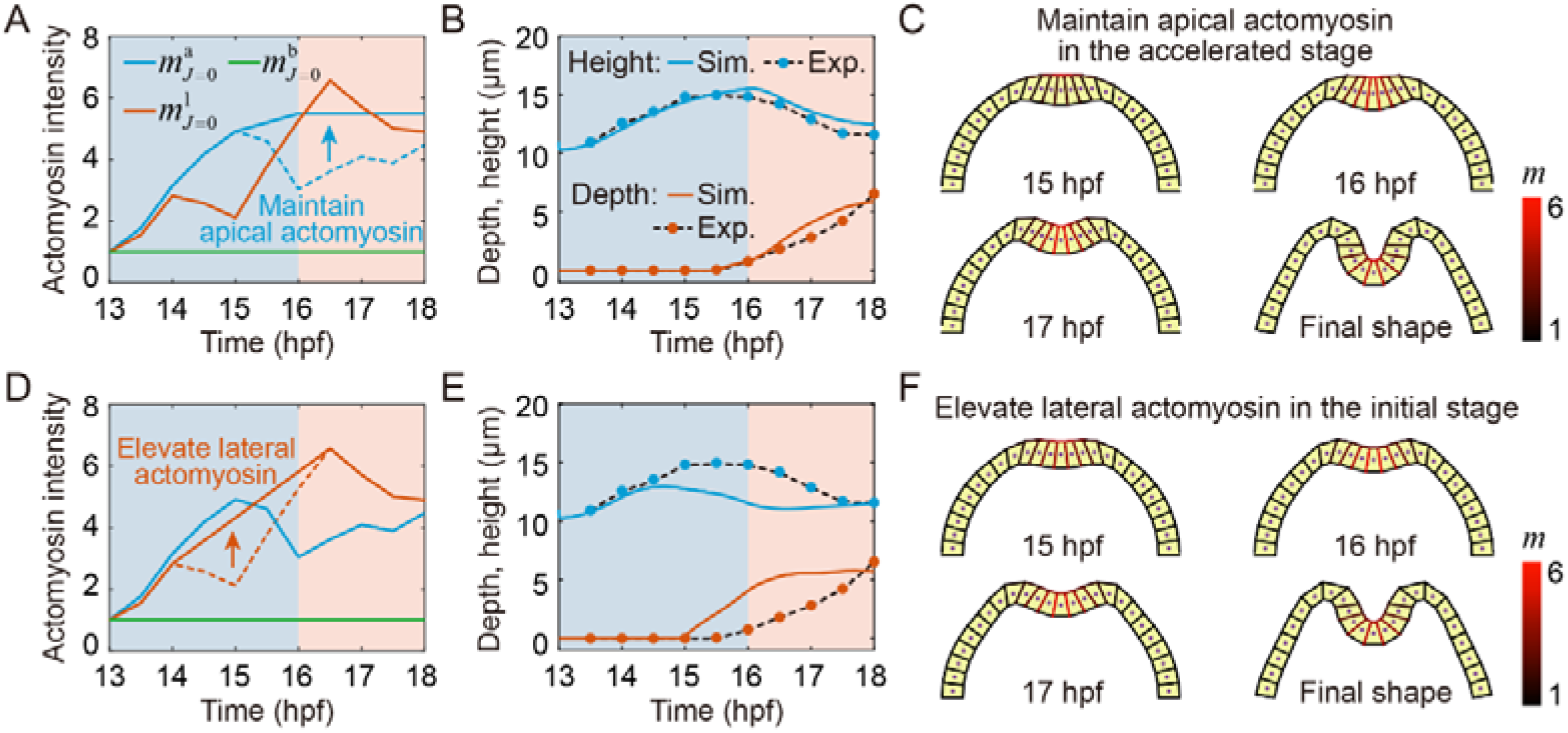
Role of actomyosin redistribution during siphon invagination. (A-C) Effects of maintaining apical actomyosin during the accelerated invagination stage. Simulations with sustained apical actomyosin (A) show the corresponding evolution of central cell height and invagination depth (B), as well as tissue morphologies at representative time points (C). (D-F) Effects of elevated lateral actomyosin during the initial stage. Simulations with elevated lateral actomyosin (D) show the corresponding evolution of central cell height and invagination depth (E), as well as tissue morphologies at representative time points (F). The dashed lines in (A) and (D) represent the experimentally measured F-actin intensities at the apical and lateral regions of the central cells, respectively.

**Appendix 1—table 1.** Key resource table.

| REAGENT or RESOURCE | SOURCE | IDENTIFIER |
| --- | --- | --- |
| antibodies |  |  |
| Phospho-Myosin Light Chain 2 (Ser19) Antibody | Cell Signaling | #3671 |
| Alexa Fluor 647 anti-Rabbit IgG | Invitrogen | #A31573 |
| Alexa Fluor 568 anti-Rabbit IgG | Invitrogen | #A11011 |
| Chemicals |  |  |
| EdU Cell Proliferation Kit with AF555 | BeyoClick | C0075S |
| Alexa Fluor™ 488 Phalloidin | Thermo fisher | A12379 |
| Protease E | sigma-Aldrich | P5147 |
| Sodium thioglycolate | sigma-Aldrich | T0632 |
| D-mannitol | sigma-Aldrich | M4125 |
| MS-222 | sigma-Aldrich | E10521 |
| VECTASHIELD® with DAPI | Vectorlabs | H1200 |
| Phanta Max Super-Fidelity DNA Polymerase | Vazyme | P505 |
| ClonExpress MultiS One Step Cloning Kit | Vazyme | C113 |
| TUNEL BrightRed Apoptosis Detection Kit | Vazyme | A113 |
| Plasmids |  |  |
| Epi1> MRLC::mCherry | This study | N/A |
| Epi1> MRLC(T18AS19A)::mCherry |  | N/A |
| Epi1> MRLC(T18ES19E)::mCherry |  | N/A |
| Epi1>PP1c::MYPT169::BcLOV4::mCherry::NES |  | N/A |
| Epi1>PP1c::MYPT169::mCherry::NES |  | N/A |
| Software |  |  |
| ImageJ | NIH Image | https://imagej.nih.gov/ij/ |
| Prism 10 CN | Graphpad Software | https://www.graphpad.com/ |
| Imaris 10 | Oxford Instruments | https://imaris.oxinst.com/ |
| MATLAB R2021a | MathWorks | https://www.mathworks.com/ |

**Appendix 1—table 2.**
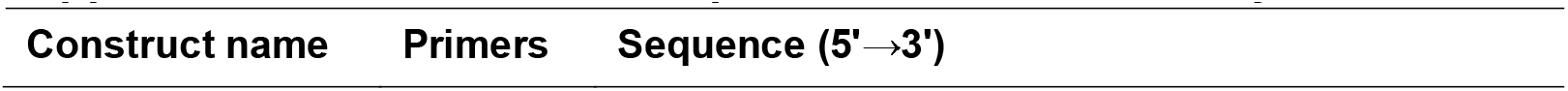
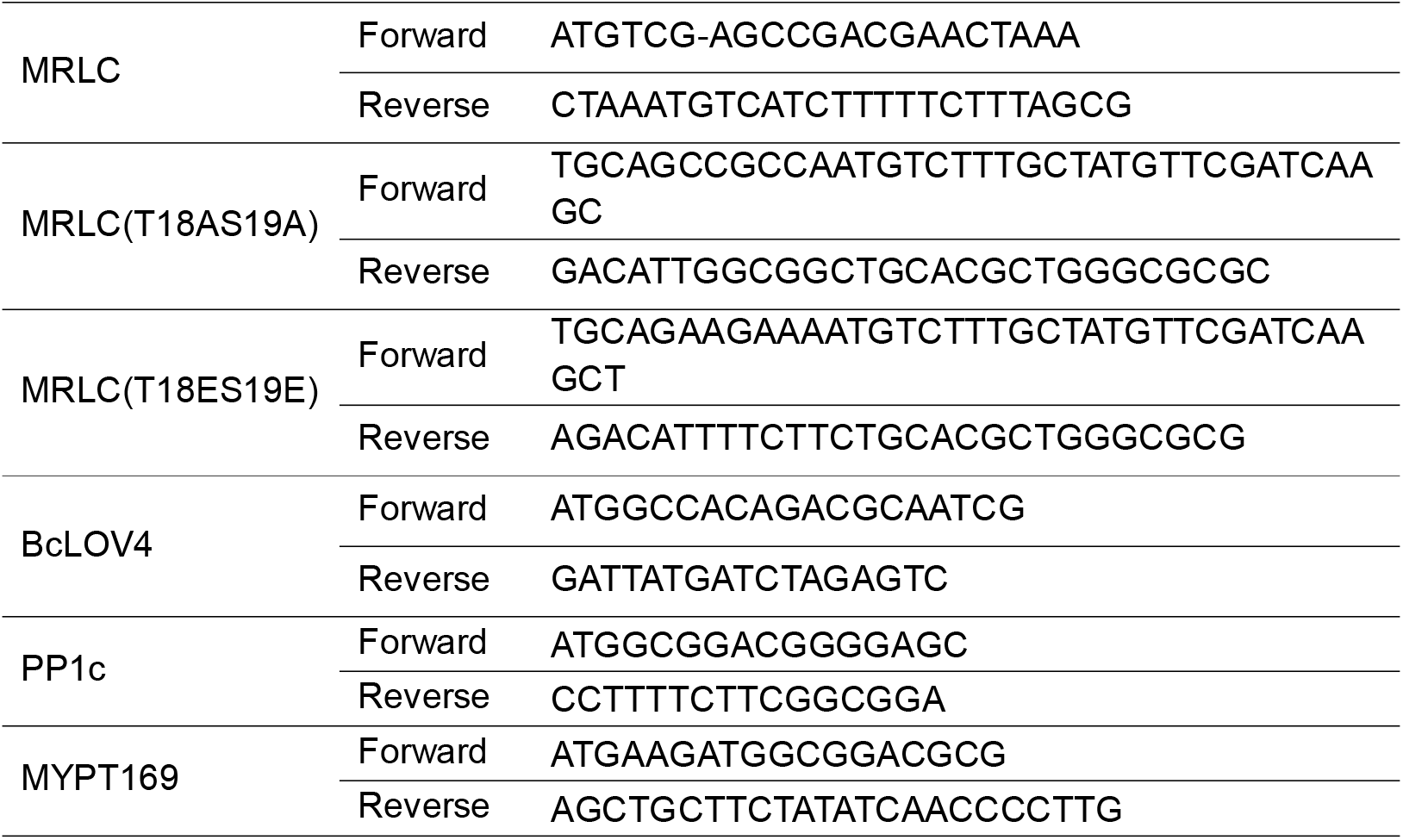
Primer sequences used in this study.

